# A Comprehensive Computational Study of Amino Acid Interactions in Membrane Proteins

**DOI:** 10.1101/617498

**Authors:** Mame Ndew Mbaye, Qingzhen Hou, Sankar Basu, Fabian Teheux, Fabrizio Pucci, Marianne Rooman

## Abstract

Transmembrane proteins play a fundamental role in a wide series of biological processes but, despite their importance, they are less studied than globular proteins, essentially because their embedding in lipid membranes hampers their experimental characterization. In this paper, we improved our understanding of their structural stability through the development of new knowledge-based energy functions describing amino acid pair interactions that prevail in the transmembrane and extramembrane regions of membrane proteins. The comparison of these potentials and those derived from globular proteins yields an objective view of the relative strength of amino acid interactions in the different protein environments, and their role in protein stabilization. Separate potentials were also derived from *α*-helical and *β*-barrel transmembrane regions to investigate possible dissimilarities. We found that, in extramembrane regions, hydrophobic residues are less frequent but interactions between aromatic and aliphatic amino acids as well as aromatic-sulfur interactions contribute more to stability. In transmembrane regions, polar residues are less abundant but interactions between residues of equal or opposite charges or non-charged polar residues as well as anion-*π* interactions appear stronger. This shows indirectly the preference of the water and lipid molecules to interact with polar and hydrophobic residues, respectively. We applied these new energy functions to predict whether a residue is located in the trans- or extramembrane region, and obtained an AUC score of 83% in cross validation, which demonstrates their accuracy. As their application is, moreover, extremely fast, they are optimal instruments for membrane protein design and large-scale investigations of membrane protein stability.

## Introduction

Biological membranes form permeable fences between the interior of cells and the external environment. They are composed of phospholipid bilayers, which form a particular, fluid, medium that differs from the surrounding aqueous solution. A lot of proteins are embedded in, attached to, or cross the membranes. We focus here on integral membrane proteins, which cross the membrane and have thus a transmembrane, an intra-cellular and an extracellular domain.

Membrane proteins are a very important class of proteins. They play key roles in the localization and organization of the cell, as well as in the cellular function by transferring specific molecules, ions and other types of signals from the cell exterior to the interior and *vice versa*. They constitute about 30% of the entire human proteome [1]. They are the focus of a lot of pharmaceutical research, as they correspond to about 60% of the current drug targets [2].

In spite of their importance, membrane proteins have been much less studied than globular proteins. They are indeed very difficult to analyze, as their folding, native structure, stability and activity is reached only within the lipid bilayer, which complicates getting their experimental X-ray structures. Generally, their large size makes also difficult to obtain them by nuclear magnetic resonance spectroscopy. These are the reasons why transmembrane protein structures only represent about 2% of the available structures deposited in the Protein Data Bank (PDB) [3]. The analysis and modeling of the 3-dimensional (3D) structure of membrane proteins are thus key objectives for rationally guiding protein design and engineering experiments.

Due to the difference between the aqueous and lipid environments, the structure and composition of transmembrane regions substantially differ from those of the intra- and extracellular domains and from globular proteins [4]. This implies that interactions that are favorable in globular regions are not necessarily so in transmembrane regions, and *vice versa*. This is a well-known fact. However, the relative strength of the different types of interactions in the two environments is not easy to evaluate.

To tackle this issue, empirical energy functions adapted to membrane proteins have been designed and used for computational modeling and design purposes (see [5, 6] for reviews). Such potentials have also been used to orient proteins into membranes, using coarse-grained molecular dynamics simulations [7, 8], or simplified potentials including anisotropic solvent models of lipid bilayers [9]. Another approach consists in deriving statistical potentials from sets of known membrane protein structures. Such potentials have been applied to evaluate structural models of membrane proteins [10, 11, 12, 13] and to position proteins into lipid membranes [10, 14].

Some authors analyzed separately *α*-helical and *β*-barrel proteins [15, 16]. Indeed, gram-negative bacteria have two membranes, an inner membrane composed of a phospholipid bilayer and an outer membrane which is an asymmetrical bilayer of phospholipids in the inner leaflet and lipopolysaccharides in the outer leaflet. This difference implies that the membrane proteins differ according to whether they are inserted in the inner or outer membrane. In particular, *α*-helical transmembrane proteins are mostly found in the cytoplasmic membranes of prokaryotic and eukaryotic cells and rarely in outer membranes, whereas *β*-barrel proteins have so far only been found in outer membranes of gram-negative bacteria, mitochondria and chloroplasts [17, 18].

In this paper, we chose to apply the statistical potential formalism to derive distance potentials from trans- and extramembrane protein regions, as this yields an objective way to compare residue-residue interactions that prevail in lipid and aqueous environments. We also derived potentials separately on *α*-helical and *β*-barrel transmembrane regions to investigate whether differences are visible between interaction strengths. We should in principle also distinguish between extramembrane residues that are in the cytoplasmic, periplasmic or extracellular regions. For example, it has been shown that positive charges in *α*-helical domains are more often situated in the cytoplasmic domain where they make interactions with the lipid molecules [19, 20], and that charged residues in *β*-proteins are more frequently located on the extracellular side [21, 22]. However, we chose to group these regions into a single category called extramembrane, which we occasionally separate into two subcategories: intracellular regions that are situated at the cellular side and are either cytoplasmic or periplasmic, and extracellular regions that can be periplasmic or really extracellular. Indeed, the number of membrane proteins with an experimental structure is currently too limited to yield reliable statistics if we define too many subregions.

## Materials and methods

### Membrane and globular protein datasets

To set up our membrane protein dataset, we used the OPM database [9], which contains experimental structures of integral membrane proteins. From these, we selected the proteins of which the structure was obtained by X-ray crystallography with a resolution of 2.5 Å at most. In a second step, we imposed a threshold on the pairwise sequence identity of 30%, with the help of the protein culling server PISCES [23]. Our final dataset 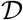 contains 165 membrane protein structures, among which 108 *α*-helical and 52 *β*-barrel polytopic integral proteins, and 5 *α*-helical monotopic integral proteins that do not span the lipid bilayer completely. They are listed in Supplementary Material Table S1.

The proteins from this dataset were divided into their transmembrane and extramembrane regions, using the OPM annotations. We got in this way two datasets, the 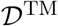 set that contains all the transmembrane protein segments, and the 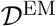 set that contains the extramembrane protein regions, and thus mix extracellular, periplasmic and cytoplasmic segments. We occasionally separated the 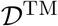 dataset into transmembrane regions with *α*-helical or *β*-barrel conformations. The protein segments that make up these datasets are specified in Table S2.

To the best of our knowledge, the dataset of protein membrane structures constructed in this paper is currently the largest non-redundant dataset used to derive effective potentials [10, 11, 12].

For comparison, we also considered the 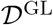 dataset set up in [24], which contains 3,823 X-ray structures of globular proteins, with a resolution of maximum 2.5 A and a pairwise sequence identity of 20 % at most.

### Statistical potentials

Statistical potentials are coarse-grained energy functions derived from frequencies of observation of associations between sequence and structure elements in a dataset of protein structures using the inverse Boltzmann law [25, 26]. In particular, we considered here the potentials:

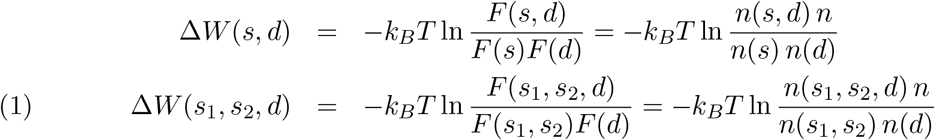

where *k_B_* is the Boltzmann constant, *T* the absolute temperature, *s, s*_1_ and *s*_2_ amino acid types, *n* numbers of occurrences and *F* relative frequencies. *d* is the spatial distance between the side chain geometric centers of two residues separated by at least one residue along the chain; the type of one of these residues (s) or of both residues (*s*_1_ and *s*_2_) are specified. The distance values between 3 and 9.9 A are divided into discrete bins of 0.3 A width and the last bin contains all distances above 9.9 A. Details about the computation of the potentials can be found in [24, 26, 27].

The potentials depend on the protein structure dataset from which the relative frequencies F are computed. Taking advantage of this dependence, a careful analysis of the relative strength of the interactions as a function of the temperature [28] and of the solubility [29] has been previously performed. Here, we extended this approach to membrane proteins and considered for that purpose the three datasets 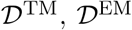 and 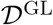. From these, we derived the transmembrane potential Δ*W*^TM^, the extramembrane potential Δ*W*^EM^ and the globular protein potential Δ*W*^GL^, which describe the interactions in these respective protein regions.

Amino acids that share similar properties can be considered together when computing the potentials. Such potentials are referred to as group potentials. In summing up the number of occurrences of different amino acid types belonging to the same group, their sizes have to be taken into account. In practice, we shifted the inter-residue distances *d* between larger amino acids towards smaller distances by subtracting the difference in radii between these amino acids and the smallest amino acid in the group. We analyzed here group potentials involving positively charged residues (Lys, Arg), negatively charged residues (Glu, Asp), aromatic residues (Phe, Tyr, Trp), aliphatic residues (Ile, Val, Leu), non-charged polar residues (Gln, Asn, Ser, Thr), small residues (Gly, Ala), and sulfur-containing residues (Cys, Met) (Table S3).

### Coping with finite-size dataset effect

Using frequencies of observation in a protein structure dataset to estimate free energy contributions through Eq. (1) implicitly assumes that the number of structures in the set is large enough to provide statistically significant values. This is, in general, a reasonable hypothesis for standard statistical potentials derived from thousands of globular structures. However, in the case of membrane proteins, the number of experimental structures is rather small and they are moreover divided into their trans- and extramembrane parts.

To cope with the finite-size effect, and get smooth and statistically significant potentials, we introduced two additional layers of computation. The first layer consists in dropping the potentials computed from distance bins *d* that do not contain a sufficient number of occurrences. We chose the threshold value on *n*(*s, d*) and *n*(*s*_1_, *s*_2_, *d*) equal to 10. If this value is not reached, the potentials are set to zero. Eq. (1) thus becomes for Δ*W*(*s*_1_, *s*_2_, *d*):

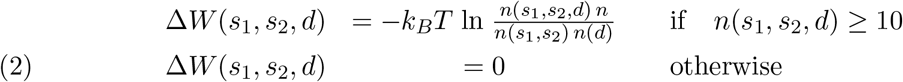

and similarly for the potential Δ*W*(*s, d*).

The second layer consists in smoothing the potential curves by replacing the number of occurrences in each bin with the weighted sum of the occurrences of the *β* neighborhood bins as:

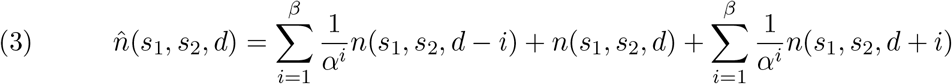

where *d* represents here a discrete distance bin rather than a continuous distance value, and where we chose *β* = 4 and *α* = 4/3. The number of occurrences 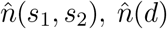 and 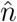 are obtained from 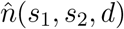 by summing over all distance bins and/or all amino acid types. The smoothing of Δ*W*(*s, d*) is done in the same way.

### Trans- and extramembrane folding free energy

The folding free energy of a protein represented by its sequence *S* and 3D conformation *C* was computed using the potentials derived from the 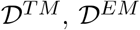 and 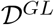 datasets as:

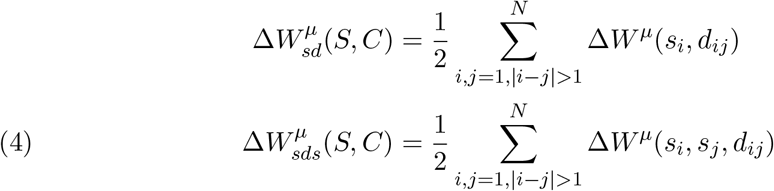

where *i, j*, and *k* denote positions along the amino acid sequence, *N* is the sequence length, and *μ* equals TM, EM or GL. To avoid any overfitting, the folding free energies were computed using a leave-one-out cross validation strategy, consisting in removing the target protein 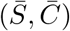 from the dataset 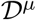 when computing its folding free energy 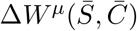. Note that this cross validation procedure is very strict, since the datasets contain, by construction, no proteins with more than 30% pairwise sequence identity.

### Per-residue folding energies

To test the accuracy and applicability of our potentials, we employed them to determine whether residues are localized in the trans- or extramembrane regions. For that purpose, we estimated the per-residue contributions to the folding free energy [30]. For residue *i*, we have:

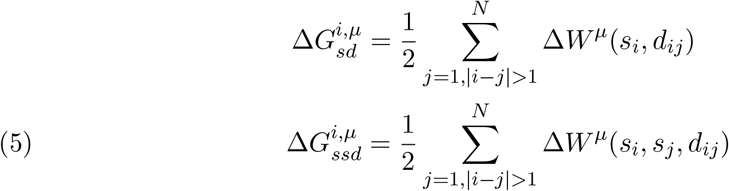

It is easy to see that the sum over all residues yields the global folding free energies of Eq. (4).

## Results and Discussion

### Amino acid frequencies

The relative frequencies of the twenty amino acids differ among the trans- and extramembrane datasets 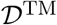 and 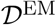, as seen in Figs 1.a and S1. Notably, the 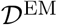 frequencies are quite similar to the 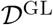 frequencies, which is not surprising as the environments of globular proteins and extramembrane regions are similar, except for the region interacting with the membrane and the transmembrane region. We also analyzed the frequency of different types of residues as a function of the distance to the intra- and extracellular water-membrane interfaces, as shown in Fig. 1.b.

**Figure 1.**
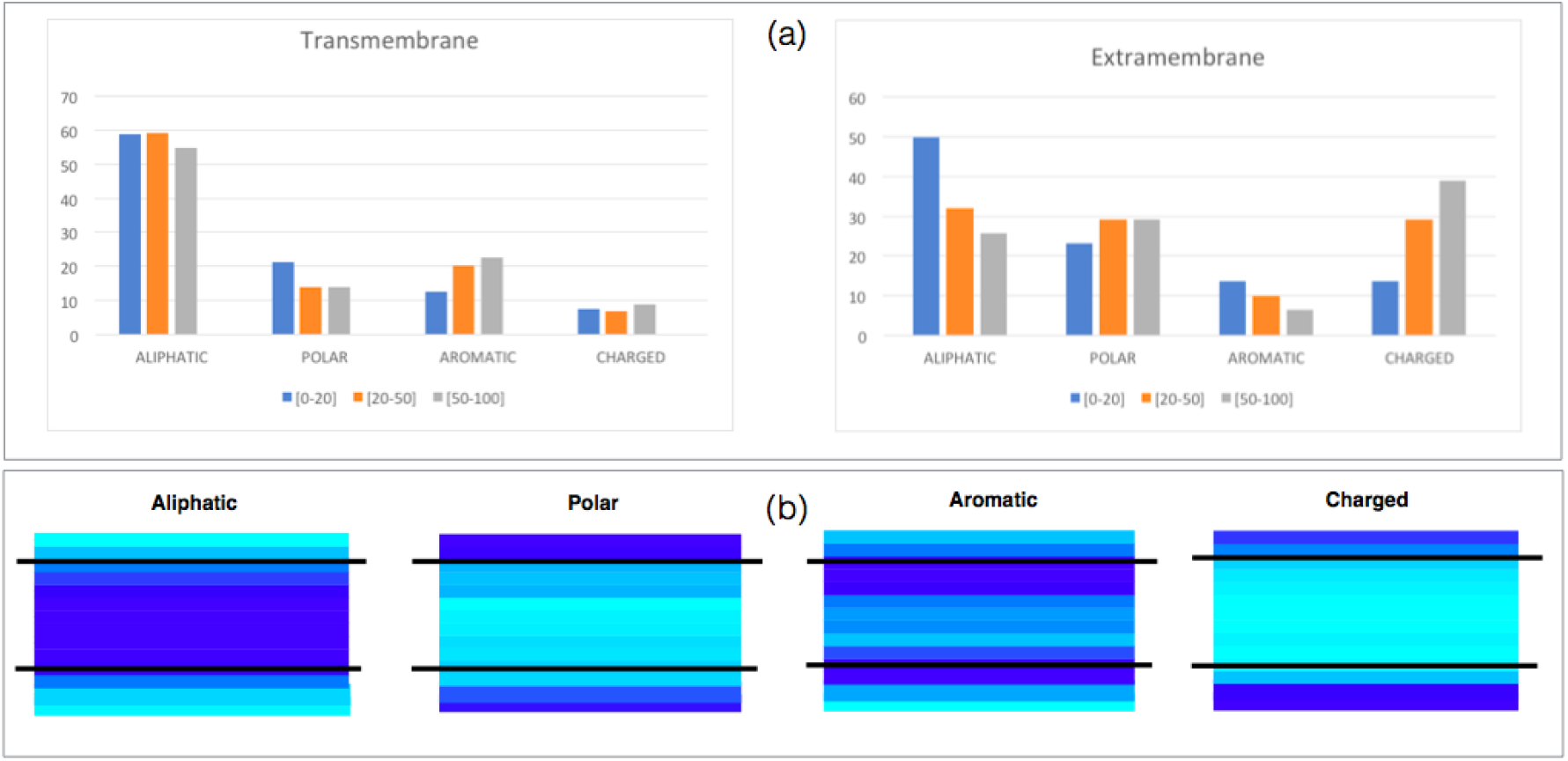
Relative frequencies of amino acid groups in the datasets 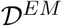 and 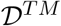. The amino acid groups are defined in Table S3. (a) Frequencies as a function of the solvent accessibility of the residues: 0-20% (blue), 20-50% (orange), and 50-100% (grey). (b) Frequencies as a function of the distance with respect to the water-membrane interfaces. Darker blue indicates higher frequencies and lighter blue lower frequencies. The straight lines represent the water-membrane interfaces. The extracellular side is directed upward and the intracellular side downward.

The clearest difference between transmembrane and extramembrane regions is observed for aliphatic residues Val, Ile and Leu: they are much more numerous in the former than in latter. In extramembrane regions, they tend to be located in the protein interior to avoid contact with water molecules, whereas in transmembrane regions, they are almost uniformly distributed; only near the interface does their frequency start to decrease. Note that Leu is more frequent than Val and Ile in transmembrane regions, probably because the former are favored in *α*-helices and the latter in *β*-strands [31] and our dataset contains more *α*-than *β*-transmembrane domains.

Aromatic amino acids were also found more frequently in the transmembrane than in the extramembrane regions. They are preferentially located near the water-membrane and the protein-membrane interfaces. This observation is consistent with the finding that aromatic residues are very important in anchoring the protein into the membrane where they tend to form cation-*π* interactions with some positively charged lipid head groups [32, 33, 34].

In contrast, charged amino acids are much more frequently observed in extra-than in transmembrane regions. This results from the large energetic cost of transferring a charged amino acid from an aqueous environment with a high dielectric constant (*ε*_water_ = 80) to the membrane that has a low dielectric constant (*ε*_membrane_ = 2 to 4) [35]. Moreover, we found differences in the distribution of positively charged residues in proteins whose transmembrane domain is *α*-helical. Indeed, as seen in Fig. S2, their frequency is higher in the regions oriented towards the cell interior than towards the cell exterior. This is consistent with the “positive-inside rule”, stating that positive residues are more abundant in the cytoplasmic regions than in the periplasmic regions for *α*-helical transmembrane domains inserted in bacterial inner membranes, or than extracellular regions in the case of eukaryotic membranes [36]. In cytochrome P450, the insertion or deletion of positively charged residues in some loop regions have been shown to modify the protein orientation with respect to the membrane and the translocation of protein segments across it [37, 38]. The general explanation of this rule is that the interaction of the positively charged residues of the intracellular domain with the negatively charged lipids of the cytosolic membrane surface through electrostatic interactions causes the retention of the positively charged residues on the cytoplasmic face of the membrane [39, 40, 41]. Note that the positive-inside-rule has been used to predict the transmembrane orientation of *α*-helical membrane proteins [42].

In *β*-barrel membrane proteins inserted into outer bacterial membranes, no significant differences are visible in Fig. S2 between charged residue frequencies in the intra- and extracellular regions. Yet, a compositional asymmetry has been described before, with a larger frequency of both positively and negatively charged residues in the extracellular regions [43, 44], where lipopolysaccharides are generally attached to the membrane. This “charge-outside” rule is not observed in our dataset.

Like the charged residues, the uncharged polar residues are also preferentially located in the extramembrane regions rather than inside the membrane. Their frequency is almost identical at both sides of the membrane.

### Preferred interactions in transmembrane regions

Statistical distance potentials were derived separately from the datasets 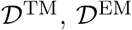 and 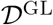, as described in Methods. Their comparison yields an objective evaluation of the residue-residue interactions that are more favorable in the transmembrane than in the extramembrane regions, and than in globular proteins. The potentials so obtained are depicted in Figs S3 and S4. In Figs S5 and S6 the potentials are computed separately for *α*-helical and *β*-barrel transmembrane regions.

#### Salt bridge interactions

Salt bridges are electrostatic interactions between positively (Lys, Arg) and negatively charged (Glu, Asp) residues which play an important role in the stabilization - especially thermostabilization [45] - of globular proteins. Here we studied the energetic contributions of this kind of interaction in the different regions of membrane proteins as a function of the distance between the residues’ side chain geometric centers. As shown in Fig. 2.a, both the extra- and transmembrane potentials have a characteristic minimum at a distance of about 4 Å^1^, but the latter are shifted downwards, by about −0.6 kcal/mol, over the whole distance range. Salt bridges appear thus much more stabilizing in the transmembrane than in the extramembrane region.

**Figure 2.**
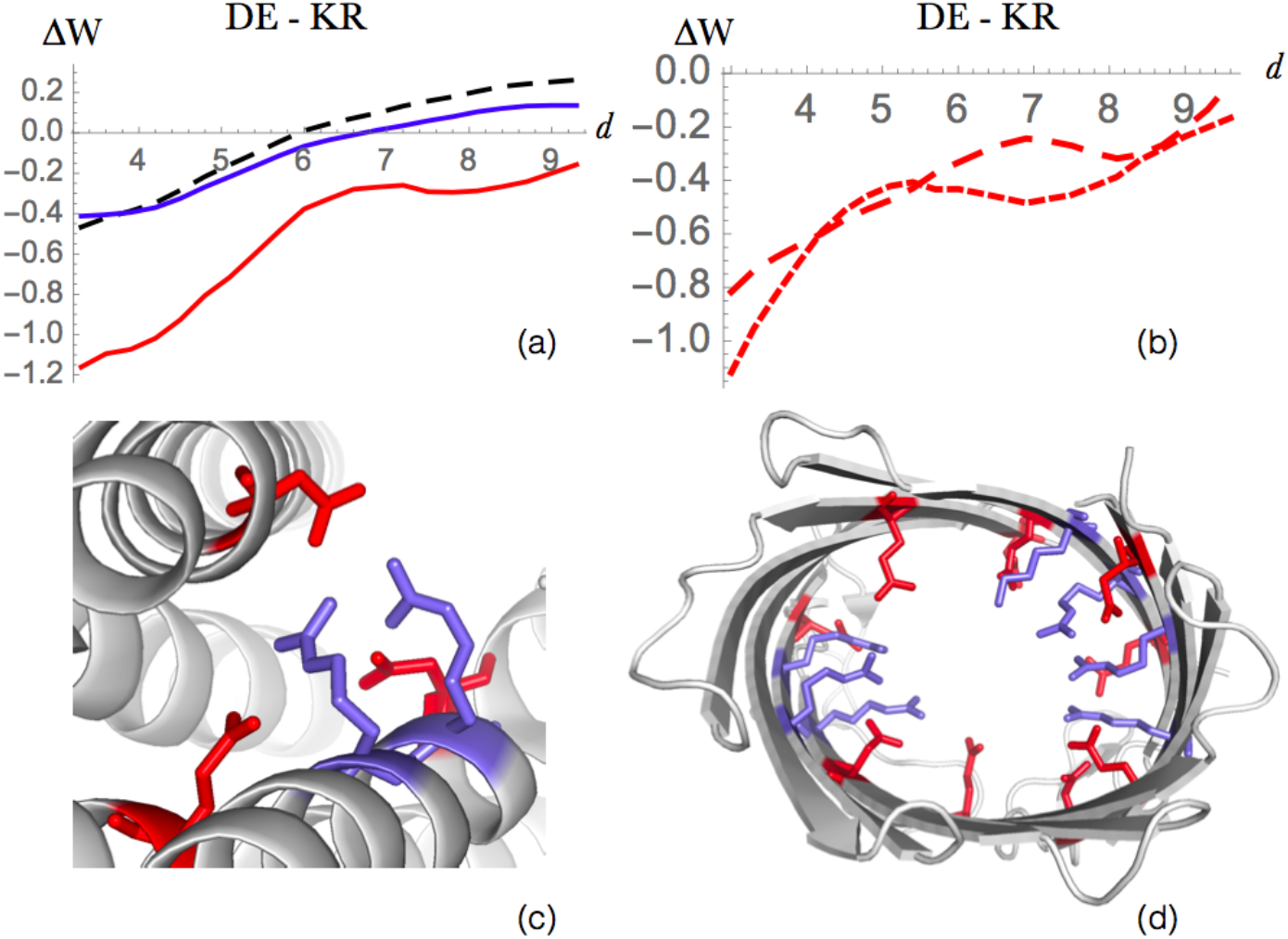
Salt bridge interactions between Arg or Lys and Asp or Glu. (a) Energy profile (in kcal/mol) as a function of the interresidue distance *d* (in Å) in globular proteins (dashed black line), extramembrane regions (blue line) and transmembrane regions (red line). (b) Difference between the energy profiles of salt bridges involving Arg (small dashed red) and Lys residues (large dashed red). (c) and (d) Salt bridges occurring in the transmembrane region of the protein structures[3] 5AYN (iron transporter ferroportin) and 2WJR (NanC porin), respectively. The residues Arg and Lys are drawn in red, and Glu and Asp in blue.

Two energy contributions play a role in the formation of salt bridges in globular proteins: the desolvation penalty upon burying an ion inside the protein, which is usually counterbalanced by the electrostatic gain in approaching the two opposite charges. In transmembrane protein regions, the situation is substantially different because the protein interior is more hydrophilic than the surface that is in contact with lipid molecules: the dielectric constant of the lipid bilayer is *ε*_membrane_ ≈ 1 – 2, whereas *ε*_interior_ varies from 2 – 6, up to 80 in the case of the hydrophilic channel in *β*-barrel porins or *α*-helical aquaporins [46]. Thus, burying an ion constitutes here an energy gain, which is added to the stabilizing electrostatic interaction between the two charged residues. We also observe that, in the transmembrane regions, Lys-containing salt bridges tend to be less stabilizing than Arg-containing ones (Fig. 2.b), in which the positive charge is delocalized on the guanidinium group.

The salt bridge geometries vary according to the type of proteins. For example, stabilizing salt bridges are recurrently found across transmembrane helices in “charge zipper” conformations, defined as extended salt bridge ladders along transmembrane helical segments [47], as illustrated in Fig. 2.c. In other membrane proteins such as porin-like *β*-barrel structures, a large network of salt bridge interactions is observed in the hydrophilic pore, as shown in Fig. 2.d.

Note that salt bridges have sometimes also pivotal functional roles. For example, they are responsible for G protein-coupled receptor (GPCR) activation and trafficking[48] and for ion channel gating [49].

#### Interactions between amino acids of equal charge

Here we focused on electrostatic interactions between two positively or two negatively charged residues, which are commonly known to be unfavorable. As seen in Fig. 3.a-b, this is indeed the case when these interactions are established between residues in globular proteins or extramembrane domains. In contrast, when two amino acids of equal charge are both in the transmembrane domain, the interaction becomes stabilizing. This can be explained by the solvation gain obtained by burying the charged residues in the more hydrophilic core or by locating them inside hydrophilic channels, which tends to dominate the repulsive electrostatic force between the two electric charges.

**Figure 3.**
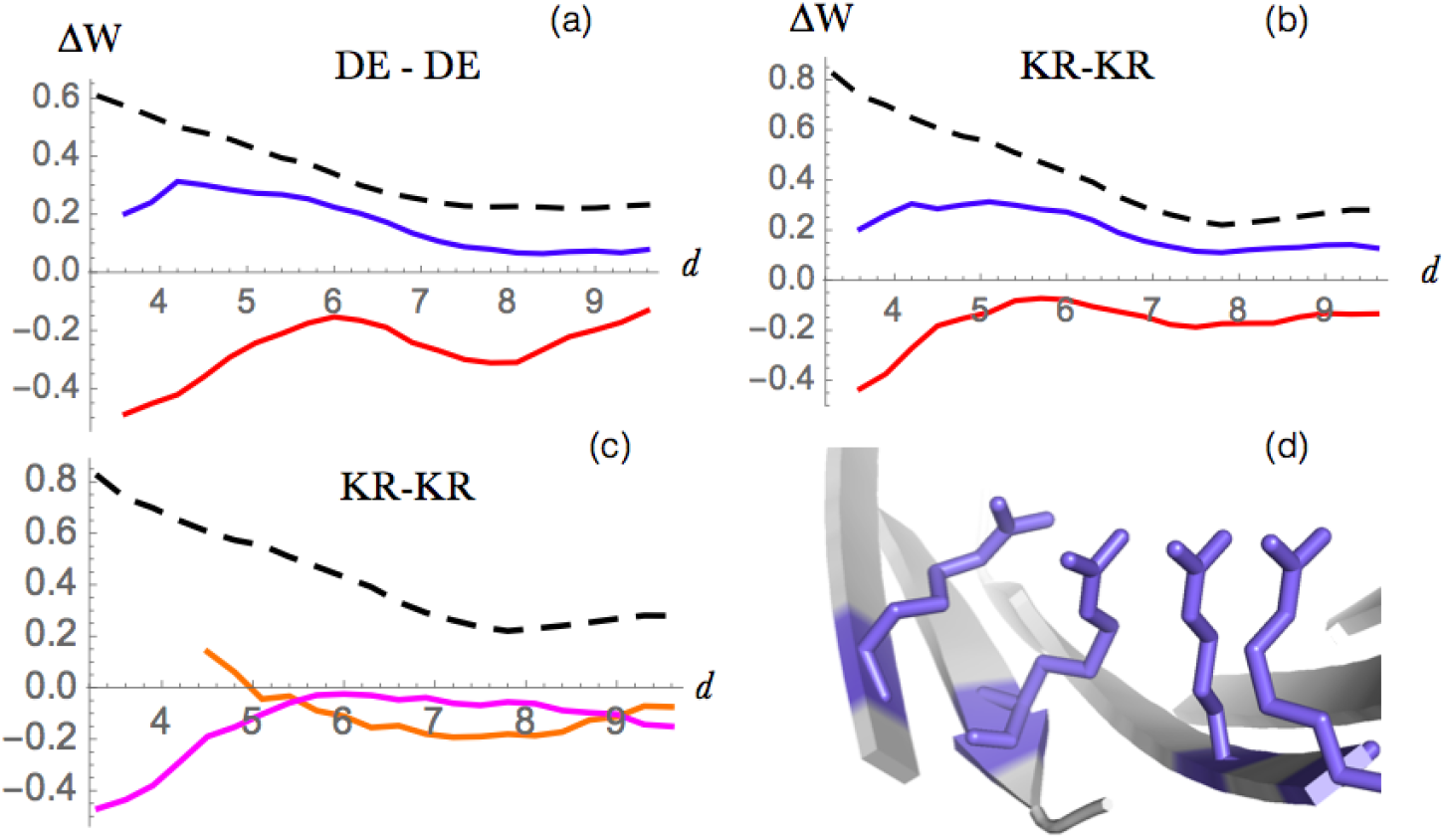
Interactions between amino acids of equal charge. Energy profiles (in kcal/mol) as a function of the interresidue distance *d* (in Å) for interactions: (a) between two negatively charged residues (Asp or Glu), and (b) between two positively charged residues (Lys or Arg), in globular proteins (dashed black line), extramembrane regions (blue line) and in transmembrane regions (red line). (c) Energy profiles of +/+ interactions in the transmembrane regions of *α*-helical proteins (orange) and *β*-barrel proteins (magenta). (d) Example of a cluster of four arginines separated by less then 4 Å inside the hydrophilic channel of the transmembrane region of KdgM porin from the *Dickeya dadantii* (PDB code 4FQE).

Surprisingly, these effective potentials become even more favorable at short distances, in spite of the electrostatic repulsion. As seen in Fig. 3.c, this counterintuitive effect is actually driven by *β*-barrel proteins, while in *α*-helical proteins +/+ and −/− interactions are very rare. Usually, we found such interactions to be located in the hydrophilic channel interior of transmembrane *β*-barrel structures. This can be explained by the earlier observation [50] of favorable clusters of positively or of negatively charged residues in interaction with water molecules. Note that this stabilizing effect is amplified for residues in which the charge is delocalized. In Arg, where the charge is delocalized on the guanidinium group, the dispersion forces between stacked guanidinium groups reduces the electrostatic repulsion. An example of an Arg cluster is given in Fig. 3.d.

#### Other polar-polar interactions

Not only the interactions between two charged residues, but also those between two non-charged polar residues, or between one charged and one non-charged polar residue, were found to be much more favorable in the transmembrane than in the extramembrane regions, and even more so, than in globular proteins (Fig. 4). The shift between the potentials is, however, smaller than for charge-charge interactions: about 0.4 kcal/mol at small distances. Note that the stabilization effect is slightly larger in *β*-barrel transmembrane proteins than in *α*-helical proteins due to the fact that the former are often channel-like structures filled with water, with which the polar moieties make favorable interactions.

**Figure 4.**
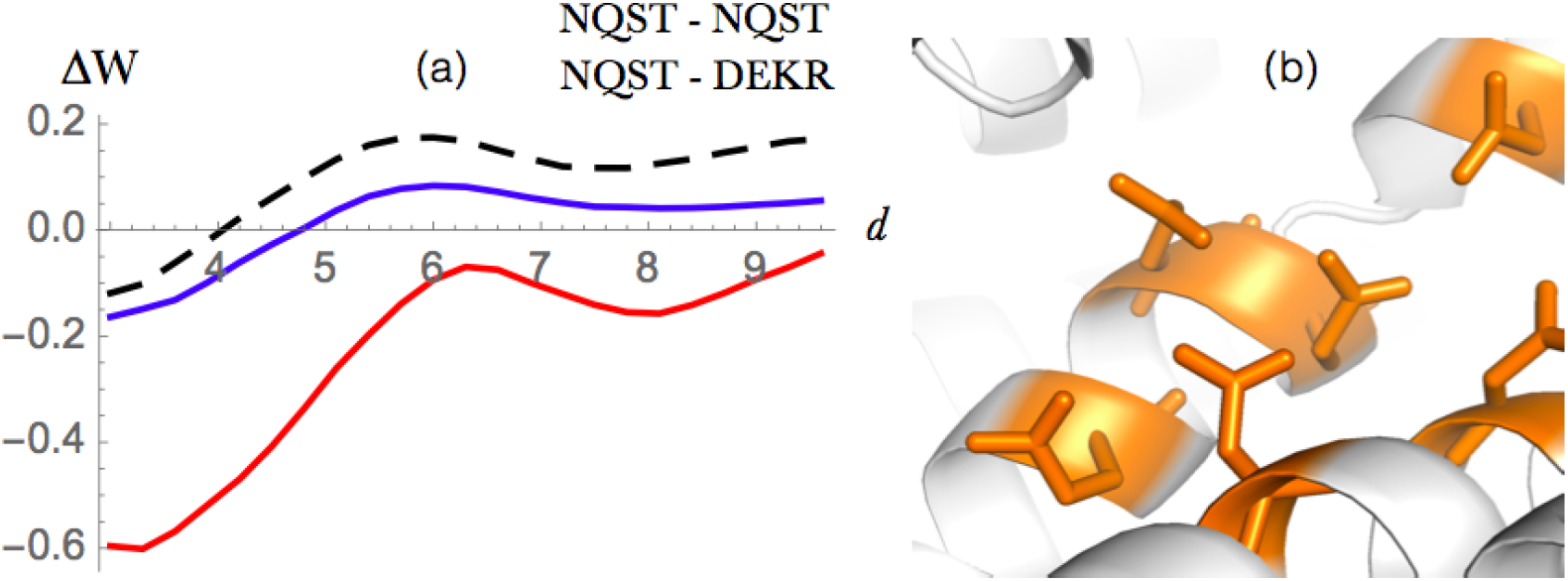
Polar-polar and polar-charge interactions. (a) Energy profile (in kcal/mol) as a function of the interresidue distance *d* (in Å) for globular proteins (dashed black), extramembrane regions (blue) and transmembrane regions (red). (b) Cluster of polar residue interactions at the interface between transmembrane helices in 4ZW9 (GLUT3 glucose transporter).

Buried polar residues have previously been described as contributing significantly to the stability of membrane protein structures [51], and to be especially important in the helix-helix interactions and in homo-oligomerization processes [52, 53]. An example of polar cluster is shown in Fig. 4.b.

#### Anion - π interactions

Since aromatic rings have non-vanishing quadruple moments, they can establish edgewise interactions with Asp and Glu side-chain carboxylate ions. Only recently has this kind of interaction received special attention in the context of their contribution to protein stabilization [54, 55, 56]. Even though some analyses suggest that their contribution is slightly destabilizing, their high occurrence frequency in biomolecular structures can be taken to signal cooperative phenomena involving other charged or aromatic residues, in which stability compensations could occur through more complex geometries such as anion-*π*-cation or anion-*π*-*π* systems [55, 56].

Fig.5 confirms that the effective energy contributions of anion-*π* interactions are destabilizing in both extramembrane regions and globular proteins, whereas their minimum value becomes neutral in the transmembrane part. Note that in the center of *β*-barrel membrane proteins, the anion-*π* interactions occur prevalently in complex geometries such as the one depicted in Fig.5.b involving two anions, two cations and two aromatic residues interacting with the aqueous solvent. In helical transmembrane regions, aromatic residues sometimes establish anion-*π* interactions with phospholipid anions; this occurs prevalently at the lipidwater interface [54].

**Figure 5.**
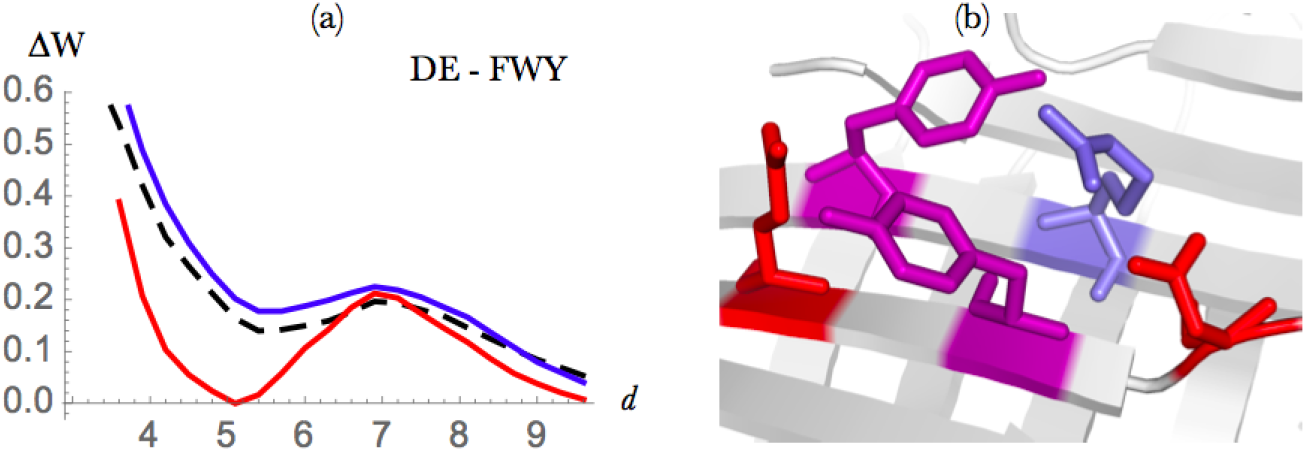
Anion-*π* interactions between Asp or Glu and Phe, Tyr or Trp. (a) Energy profile (in kcal/mol) as a function of the interresidue distance *d* (in Å) in globular proteins (dashed black), extramembrane regions (blue) and transmembrane regions (red). (b) Example of an anion-*π* interactions in the PDB structure 1A0S (sucrose-specific porin). The negatively charged amino acids are in red, the positively charged ones in blue and the aromatic residues in magenta.

#### Cation-πinteractions

Cation-*π* interactions are established when the cationic side chain of Lys or Arg is localized above or below the aromatic ring of Phe, Trp or Tyr. They play an important role in the stabilization of protein structures of both membrane and globular proteins and in protein-protein, protein-DNA and protein-ligand complexes [57, 58, 59, 60, 61].

The distance-dependent energy profile of this kind of interactions is depicted in Fig. 6.a. The potentials extracted from transmembrane, extramembrane and globular regions are similar, with a slightly more negative curve at short distances (<4 Å) in the case of globular proteins, and a preference for transmembrane regions with respect to the extramembrane ones for <6 Å.

**Figure 6.**
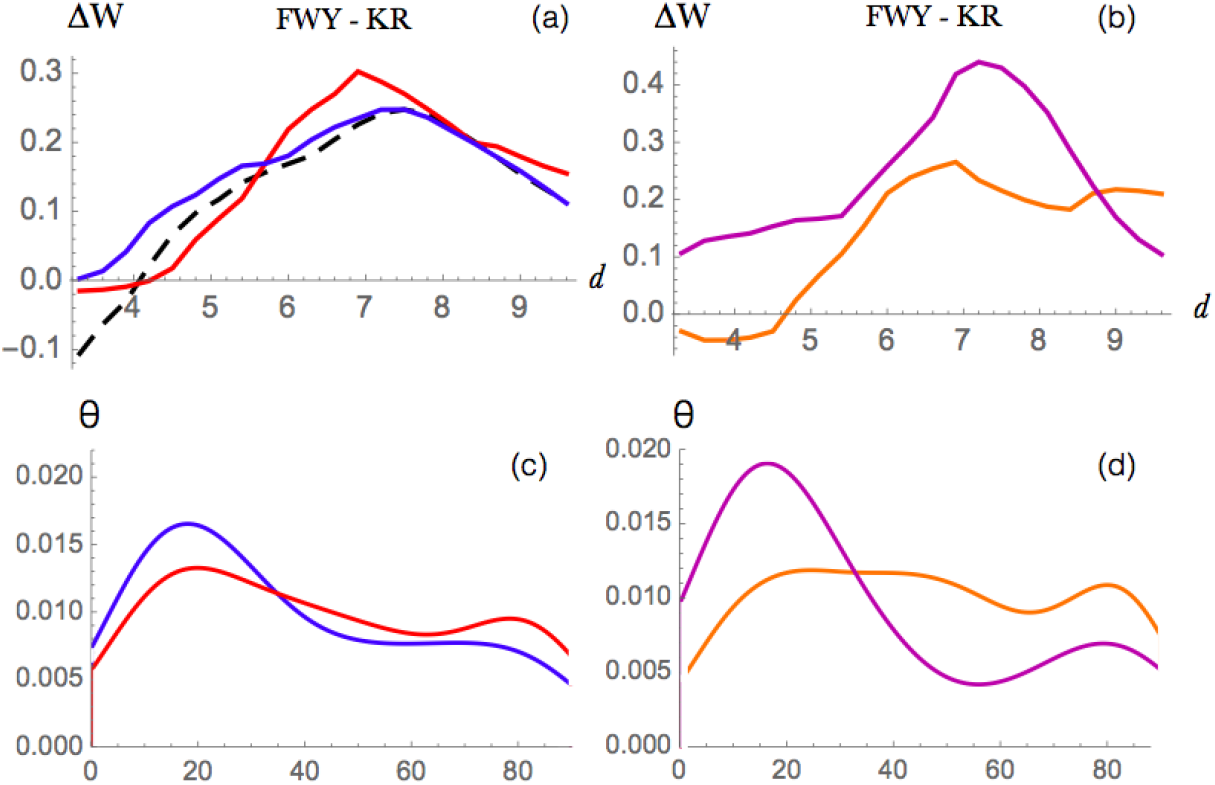
Cation-*π* interactions between Lys or Arg and Phe, Tyr or Trp (a) Energy profile (in kcal/mol) as a function of the interresidue distance *d* (in Å) in globular proteins (dashed black), extramembrane regions (blue) and transmembrane regions (red). (b) Energy profile in *α*-helical (orange) and *β*-barrel (magenta) transmembrane regions. (c)-(d) Distribution of the *θ* angle between the aromatic and guanidinium planes, for Arg-involving cation-*π* interactions. 0° corresponds to stacked and 90° to parallel conformations. (c) Distributions from extramembrane regions are in blue and those from transmembrane regions in red. (d) Distributions from *α*-helical (orange) and *β*-barrel (magenta) transmembrane proteins.

It has been suggested that cation-*π* interactions influence more strongly *β*-barrel than *α*-helical transmembrane proteins [62]. In order to objectively study this difference, we plotted cation-*π* energy profiles extracted from these two different protein classes (Fig.6.b). What we found differs from previous findings [62]: the energy profile at short distances (below 5 Å) is negative in *α*-helical and slightly positive in *β* proteins. This indicates that cation-*π* interactions contribute more to stability in *α*-helical transmembrane regions.

In cation-*π* interactions involving Arg, the planar guanidinium group and the aromatic moieties can make favorable stacking interactions, which add up to the electrostatic interactions. We analyzed the geometry of these interactions through the study the distribution of the angle between the aromatic and guanidinium planes. As shown in Fig. 6.c-d, the angle is preferentially around 20° in *β*-barrel transmembrane regions and the two planes are thus almost in parallel, stacked, conformations. In extramembrane regions, a preference for stacked conformations is also visible, whereas in *α*-helical transmembrane regions, basically all angle values are observed.

Cation-*π* interactions are known to be important not only for stability but also for their functional roles such as for example in substrate and ligand binding [63, 61]. When they are established between the aromatic residues of the protein and the positively charged portion of phospholipid head groups, they are fundamental to anchor the protein to the membrane [32, 33, 34]. The importance of the aromatic rings in membrane anchoring is not easy to show using the statistical potential formalism as the so-obtained effective potentials take only implicitly the impact of the environment into account; indications of this anchoring effect are observed from the aromatic amino acid frequencies in Fig 1.

### Preferred interactions in extramembrane regions

We now have a closer look at the residue-residue interactions that are more favorable in the extramembrane than in the transmembrane regions, as measured by the distance potentials.

#### Sulfur-aromatic interactions

Sulfur-containing amino acids (Cys and Met) are highly polarizable and can establish nonbonded interactions with aromatic moieties. It has been shown that they play important roles not only in the stabilization of protein structures [64, 65, 66, 67] but also in their function [67, 68], as for example in the protection of Met against oxidation leading to methionine sulfide.

The potentials in Fig. 7.a show the stabilizing contribution of sulfur-aromatic interactions, which is much stronger for the extramembrane than for the transmembrane regions. Indeed, for the latter region, the entire energy profile is shifted by about +0.2 kcal/mol on the average over all distances. It is interesting to note that sulfur-*π* interactions in transmembrane regions occur almost exclusively in *α*-helical proteins where interhelical interactions frequently involve methionine surrounded by a cage of aromatic residues. In the extramembrane region, they frequently involve partially exposed residues and more sulfur than aromatic residues (Fig. 7.C).

**Figure 7.**
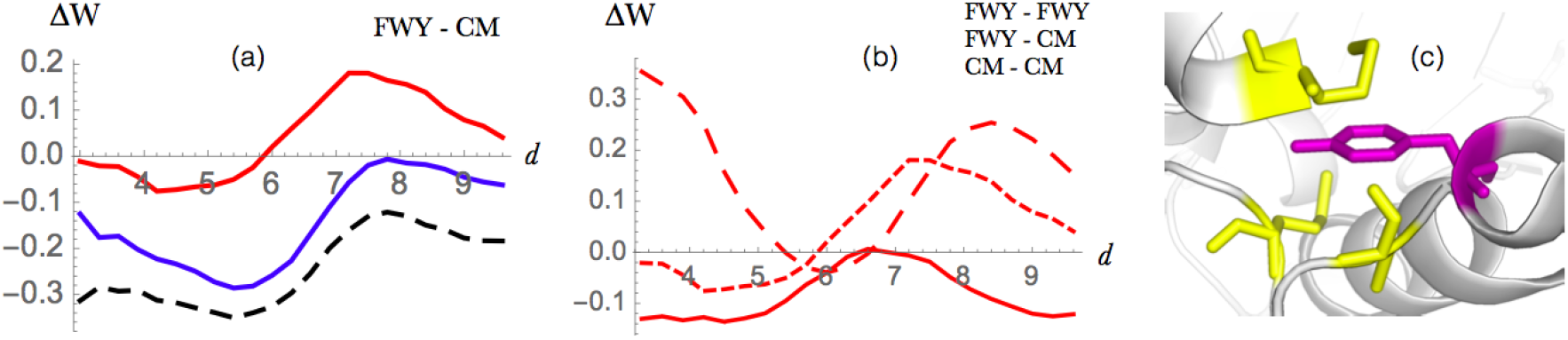
Sulfur-*π* interactions between Met or Cys and Phe, Tyr or Trp. (a) Energy profile (in kcal/mol) as a function of the interresidue distance *d* (in Å) in globular proteins (dashed black), extramembrane (blue) and in the transmembrane regions (red). (b) Energy profile for sulfur-*π* (small dashed red), *π*-*π* (large dashed red) and sulfur-sulfur (continuum red) interactions in the transmembrane regions. (c) Example of sulfur-*π* interaction in the transmembrane region of the PDB structure 3S8G (ba3 cytochrome c oxidase); Met and Cys are in yellow and aromatic residues in magenta.

We compared the strength of sulfur-*π* interactions and of aromatic-aromatic and sulfur-sulfur interactions in the transmembrane regions, but did not find a clear difference between the minimum energy values (Fig 7.b). This contrasts with earlier results obtained by a combination of structural bioinformatics and *ab initio* quantum chemistry calculations, which suggested that sulfur-aromatic interactions in membrane proteins are more stabilizing than aromatic-aromatic or sulfur-sulfur interactions [66].

Regarding the geometry of the sulfur-*π* interactions, we did not see any substantial difference between the trans- and extramembrane regions. In both regions, we observed a slight preference for conformations with an angle of about 40-45° between the sulfur and the normal vector defined by the plane of the aromatic ring, in agreement with earlier findings [64].

#### Aromatic interactions

Due to their hydrophobic nature, especially marked for Phe, aromatic amino acids prefer to be located in transmembrane regions or in the core of extramembrane regions (Fig. 1). On the basis of their energy profiles (Fig. 8.a), we observed that the interactions between pairs of aromatic residues are more favorable in extramembrane than in transmembrane regions. Moreover, they have almost the same weight in *α*-helical and *β*-barrel proteins, with a slight preference for the former (Fig. 8.b), in agreement with earlier studies [69]. Note that in *β*-barrel proteins, the aromatic residues are usually lipid-facing, whereas in *α*-helical proteins they are in the protein interior. This difference is due to the fact that *β*-barrel transmembrane regions have almost no core.

**Figure 8.**
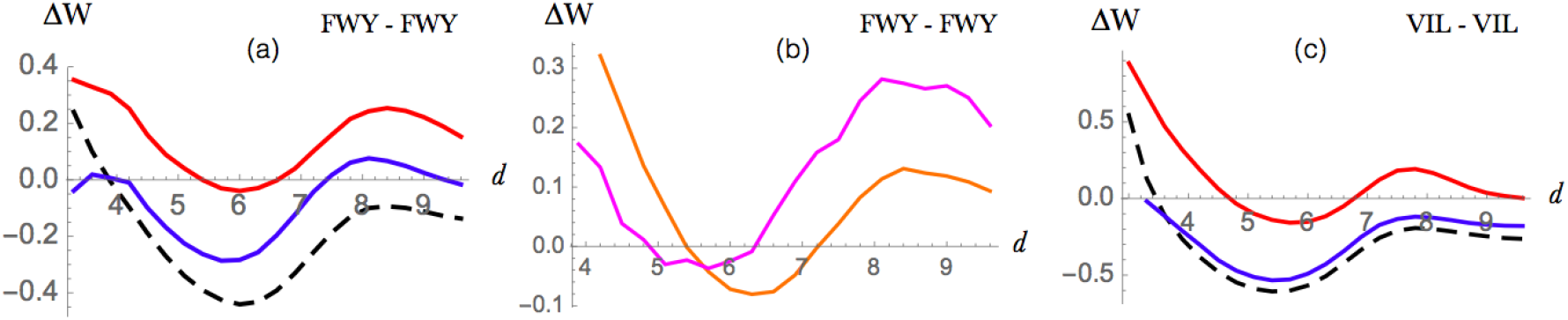
Aromatic-aromatic interaction between Phe, Tyr or Trp and aliphatic-aliphatic interactions between Ile, Leu or Val (a) Energy profile (in kcal/mol) of aromatic-aromatic interactions as a function of the interresidue distance *d* (in Å) in globular proteins (dashed black), extramembrane (blue) and transmembrane regions (red). (b) Energy profile for aromatic-aromatic interactions in *α*-helical proteins (orange) and in *β*-proteins (magenta). (c) Energy profile (in kcal/mol) for aliphatic-aliphatic interactions as a function of the distance *d* (in Å) in globular proteins (dashed black), extramembrane (blue) and transmembrane regions (red).x

The geometries of the aromatic-aromatic interactions are similar between those occurring in transmembrane and extramembrane regions (data not shown). They occur preferentially in a T-shaped conformation. Note that *π*-*π* stacking plays a role not only in the tertiary structure stabilization but also in the oligomerization of the membrane protein subunits [69].

When aromatic amino acids are positioned close to the lipid interface, they are known to play important roles in anchoring and positioning the protein inside the lipid medium through lipid-aromatic interactions (see [70, 71]). The interactions between amino acids and lipid molecules are, however, not captured by our statistical potentials, which consider both lipids and water as the protein environment.

#### Aliphatic interactions

While hydrophobic forces play a dominant role for folding and stability in globular proteins, they contribute less to the stability of the transmembrane proteins [72]. This is indeed exactly what we observe in the energy profiles of Fig. 8.c. When the interactions are established in extramembrane regions, the potentials are clearly stabilizing with an energy minimum at about 6 Å like in globular proteins. In transmembrane regions, the minimum is still present but about 0.4 kcal/mol higher, which indicates that these interactions are only marginally stabilizing.

However, even though hydrophobic forces are less important for folding, they are one of the contributing factors for the positioning and anchoring of the protein to the lipid membrane, especially in peripheral membrane proteins [72] but also in integral membrane proteins. Indeed, hydrophobic interactions can be established between exposed non-polar residues and hydrophobic lipid moieties of the membrane, which determine the insertion and position of the proteins [73]. There are indeed more and more indications of protein-membrane hydrophobic matching, in which the hydrophobic part of the transmembrane domain has to match the hydrophobic thickness of the membrane bilayer in which it is embedded; moreover, this matching condition appears to strongly influence protein function [73]. Since our statistical potentials take implicitly but not explicitly the membrane bilayer into account, the latter effects are only observed indirectly.

### Application of the membrane potentials to predict residue localization

The newly developed membrane statistical potentials were used to perform a binary classification of the residues into those that belong to the transmembrane or extramembrane regions. We computed for that purpose the per-residue contributions to the folding free energy derived from the extra- and transmembrane datasets 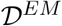 and 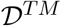 defined in Eq. (5). In general, we expect that if the per-residue contribution computed with transmembrane potentials is lower than that computed with extramembrane potentials, the residue in situated inside the membrane, and *vice versa*. But there are sometimes deviations from this rule. Indeed, some residues correspond to stability weaknesses, which means that they contribute unfavorably to the overall folding free energy [74, 30].

To predict the localization of a residue, we considered linear combinations of the per-residue folding free energies computed with the potentials “sd” and “sds” from the two datasets 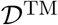 and 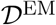:

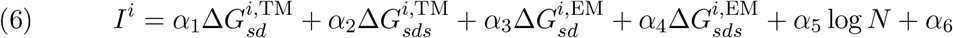

where the coefficients *α* are parameters. We added two terms in this localization index: a constant term and a term proportional to the logarithm of the protein length. The latter term is introduced to correct for the possible length dependence of amino acid and distance frequencies [75]. We also defined a smoothed version of this localization index, by averaging it over a window of five successive residues along the chain centered around the target residue:

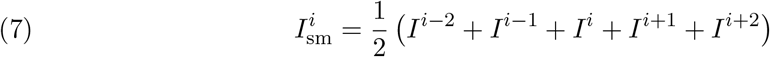

This index was used to classify the residues into two groups: the residues *i* with 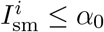 were considered to belong to the transmembrane region and those with 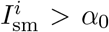 to the extramembrane region. The seven parameters *α_j_* (with *j*=0…6) were identified so as to optimize the values of the balanced accuracy (BACC); the area under the receiver operating characteristic curve (AUC) was also computed.

The tests of performance were done using a strict leave-one-out cross validation procedure, where the target protein, whose residues we want to classify, is removed in all the stages of the computations, from the derivation of the statistical potentials to the optimization of the parameters. As the pairwise sequence identity inside the datasets is low (< 30%), the cross validation is strict and in principle free from biases.

As shown in Table 1 and Fig. 9.a, we obtained a BACC of 0.75 and an AUC of 0.83 on the whole set of membrane proteins. These good results indicate that our potentials describe quite well the stability properties of the membrane proteins in the two completely different environments that are water and lipids, and thus that they can be used to localize residues inside or outside the membrane.

**Figure 9.**
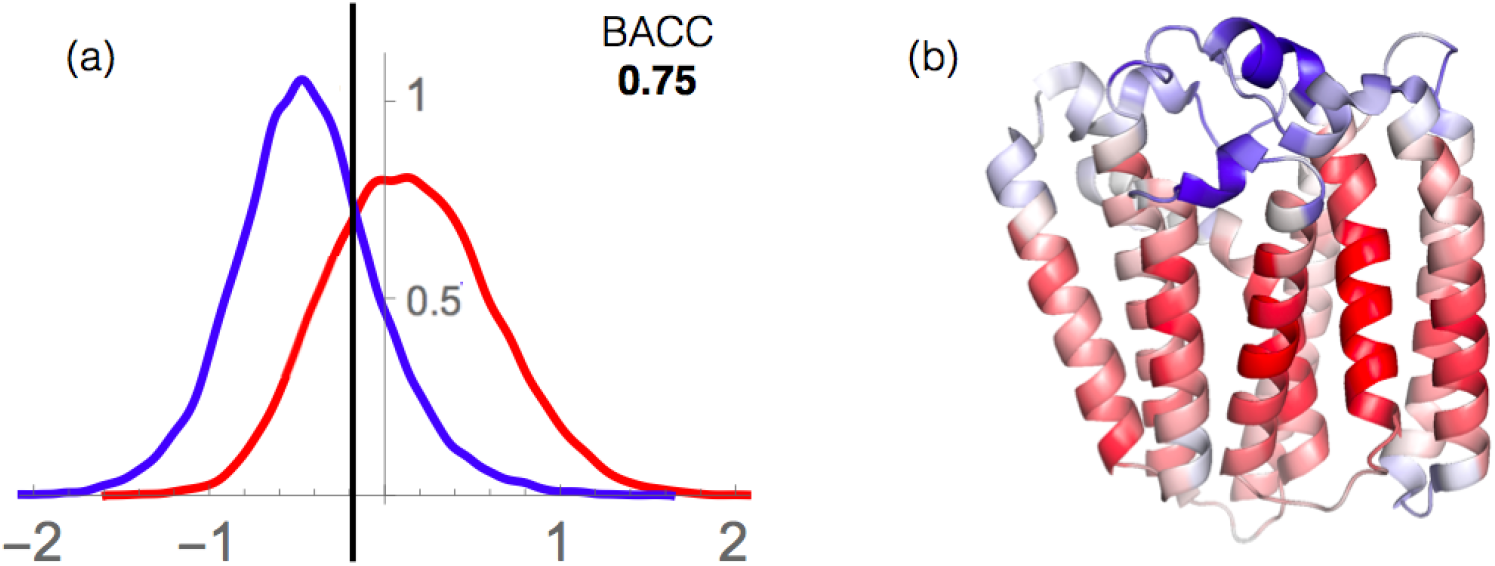
Residue localization inside or outside the membrane. (a) Distribution of the 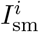 index for the transmembrane (red) and extramembrane (blue) regions. The binary classification is performed using the threshold value indicated by the vertical black line, which yields a BACC value of 0.76. (b) Representation of the 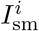 values for a prenyltransferase integral transmembrane protein (PDB code 4TQ4). The color scale, from red to blue, represents the 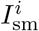 values; red indicates a strong prediction of transmembrane localization and blue, of extramembrane localization.

**Table 1.** BACC and AUC values of the prediction of residue localization (inside or outside the membrane) obtained from the index *I_sm_*. The values in parentheses indicate the number of proteins in each set.

|  | BACC | AUC |
| --- | --- | --- |
| All (164) | 0.75 | 0.83 |
| α-proteins (112) | 0.78 | 0.86 |
| β-barrels (52) | 0.67 | 0.74 |

Our predictor works better for the *α*-helical proteins (AUC=0.86) than for the *β*-barrels (AUC=0.74). We can argue that this difference is due to the fact that our dataset is dominated for two thirds by *α*-helical proteins, and that it is thus normal that this type of proteins is better predicted than *β*-barrels. Moreover, the *β*-barrel subset consist of channel and porin structures, in which the transmembrane region has an internal hydrophilic region in contact with water, and this makes this set substantially more difficult to predict using distance potentials only.

An example of localization prediction is shown in Fig. (9).b for *Archaeoglobus fulgidus* prenyltransferase, an *α*-helical integral membrane protein. Its residues are colored according to the predicted values of the localization index 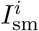. Clearly, our potentials are able to discriminate between extra- and transmembrane regions. Note that some of the residues that are not localized correctly are close to the membrane-water interface, where our potentials are the least accurate (see Conclusion). Some others could correspond to stability weaknesses, which means that they would benefit from being mutated to improve the global protein stability.

## Conclusion

In this paper, we developed new transmembrane and extramembrane residue-residue potentials in view of identifying the amino acid interactions that contribute more strongly to the stabilization of either the transmembrane or the extramembrane region, and we compared them with their interaction strength in globular proteins. First of all, we observed that the potentials derived from globular proteins are much more similar to those derived from extramembrane than from transmembrane regions.

Despite their low occurrence in transmembrane regions, it seems that interactions involving polar residues tend to contribute more to the stability of these regions than of the extramembrane regions. In particular, salt bridges are stabler by more than 0.5 kcal/mol, and interactions between residues of equal charge, which are usually destabilizing, become stabilizing when located inside the membrane. This effect can be explained by the fact that burying a charged residue inside the lipid environment is not associated with a desolvation penalty, as it is in an aqueous environment. Note that clusters of positively or negatively charged residues situated inside *β*-barrel porin channels may have, not only a structural, but also a functional role in the flux of targeted molecules through the membrane. Non-charged polar-polar and anion-*π* interactions appear also more favorable in the transmembrane region, and so do cation-*π* interactions but to a much smaller extent.

Opposite trends are observed in the extramembrane regions. Hydrophobic residues, despite their preferential location in transmembrane regions, establish stronger effective interactions in extramembrane regions due to their pronounced tendency to avoid contact with water molecules, but not with lipids. In particular, aromatic-aromatic, aliphatic-aliphatic and aromatic-sulfur interactions appear to contribute more to stability in extramemembrane regions.

Note that these results have to be understood in the context of statistical mean-force potentials in which the water and lipid molecules are not considered explicitly. The lack of interactions between polar residues in extramembrane regions is indeed counterbalanced by interactions between polar residues and water molecules. Similarly, the lack of interactions between hydrophobic residues in intramembrane regions is counterbalanced by interactions between hydrophobic residues and lipid molecules.

Moreover, the class of transmembrane proteins strongly influences the effective strength of some of the residue-residue interactions. Indeed, we observed marked differences between some potentials derived from *α*-helical and *β*-barrel transmembrane domains. This is related to the fact that the latter are all channel-like structures filled with water and that the residues pointing towards the channel interior are mostly hydrophilic, whereas only a small fraction of the *α*-helical transmembrane proteins have such a structure. In fact, *β*-barrel transmembrane regions have no real core. Another difference between these two protein classes is due to the fact that *β*-barrel membrane proteins tend to be located in the outer membrane whose characteristics differ from the internal membrane where the *α*-helical proteins are almost exclusively located. The effect of two different environments of course influences the shape of our membrane-protein statistical potentials.

In order to check the validity of our statistical potentials, we used them to predict whether a residue is localized in the transmembrane or in the extramembrane region. The high BACC and AUC values obtained in cross validation, in addition to the fact that their application is extremely fast, make these potentials an invaluable asset for various investigations in membrane protein design or in large-scale studies of membrane positioning.

Despite the good results obtained, our potentials can still be improved. Obviously, when larger larger datasets of membrane proteins will be available, our statistical potentials will certainly yield a more accurate description of the stabilizing contributions, and will make it possible to divide the dataset into several subclasses of transmembrane proteins that have specific characteristics such as ion channels, (aqua)porins, *α*-helical or *β*-barrel topology, or their insertion into different membrane types, which are likely to influence the effective interactions. Moreover, potentials that involve other structural descriptors than the interresidue distance, such as backbone torsion angle domains or solvent accessibility could further improve the prediction of residue localization presented here. This will be the subject of a forthcoming paper.

Finally, the interactions that prevail at the water-lipid or protein-lipid interface are crucial for the anchoring of transmembrane proteins into the membrane and are not well described by our statistical potentials. These are by definition effective potentials and thus the interactions with the lipid or aqueous environment are only considered indirectly. Combining the present analysis with explicit solvent models could be a possibility to unravel this important aspect of membrane proteins.

## Supporting information

**Table S1.** Membrane protein dataset.

**Table S2.** Transmembrane protein segments in the dataset.

**Table S3.** Amino acid groups.

**Figure S1.** Relative frequencies of the 20 amino acids in the datasets in the datasets 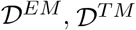 and 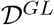 as a function of the solvent accessibility of the residues.

**Figure S2.** Relative frequencies of negatively and positively charged residues in the datasets 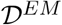 and 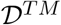 as a function of the distance with respect to the water-membrane interfaces.

**Figure S3.** Statistical sds residue-residue potentials as a function of the distance, derived from the datasets 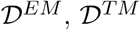 and 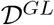.

**Figure S4.** Statistical sds potentials between amino acid groups as a function of the distance, derived from the datasets 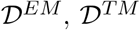 and 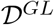.

**Figure S5.** Statistical sds residue-residue potentials as a function of the distance, derived from the dataset 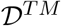, or separately from *α*-helical and *β*-barrel transmembrane regions.

**Figure S6.** Statistical sds potentials between amino acid groups as a function of the distance, derived from the dataset 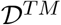, or separately from *α*-helical and *β*-barrel transmembrane regions.

## Acknowledgments

This work is supported by the FNRS Fund for Scientific Research through a PDR grant. M.N.M has a PhD grant from the Belgian Commission for Cooperation and Development (ARES-CCD); Q.H., S.B. are Postdoctoral Researchers and M.R. is Research Director at the FNRS. F.P. has been supported by the FNRS and by the John von Neumann Institute for Computing (NIC)

## Conflict of Interest

We declare that there is no conflict of interest regarding the publication of this manuscript.

## Footnotes

1 Note that this distance is rescaled towards the smallest amino acid as explained in Methods.

